# Porcine Left Atrial and Ventricular Thick Filaments Exhibit Distinct Resting Structures and Calcium-dependent Responses

**DOI:** 10.64898/2026.05.18.726029

**Authors:** Lin Qi, Maicon Landim-Vieira, Hailey Flannagan, Marcela Monroy, Edward O. Olaniyan, Meihua Guo, Chengqian Gao, Shengyao Yuan, Henry Gong, Suman Nag, Thomas C. Irving, Weikang Ma

## Abstract

The heart maintains systemic perfusion through the coordinated function of its four chambers: the left and right atria and ventricles. Each chamber has distinct structural, functional, and molecular properties tailored to its role in circulation, which may result in chamber-specific differences in myofilament structure and regulation between atria and ventricles. To test this hypothesis, we employed muscle mechanics and X-ray diffraction to investigate functional and structural differences in porcine left atrial (LA) and left ventricular (LV) tissue. Here, we report the first X-ray diffraction study of atrial tissue, demonstrating that under resting conditions, myosin filaments in LA adopted a more ON-like, structurally distinct configuration compared with those in LV. Under contracting conditions, LV generated greater force and exhibited higher sinusoidal stiffness than LA across multiple calcium concentrations. LA showed faster *k*_TR_ than in LV, with no calcium-dependence, in contrast to the calcium-dependence of *k*_TR_ seen in LV. Structurally, the distinct myosin head configuration seen in the relaxed LA persisted during contraction. Furthermore, using the troponin inhibitor MYK-7660 to inhibit active contraction, we showed that, unlike LV, LA showed no direct calcium-dependent thick filament activation, reconciling discrepancies between fast rat and slow porcine ventricular myocardium regarding calcium’s role in thick filament regulation. Altogether, our study reveals that LA myosin filaments adopt a molecular architecture and regulatory mechanism distinct from their LV counterparts, suggesting that myosin filament structure and regulation have evolved differently to meet the unique functional demands of each cardiac chamber. Moreover, atrial disease is often associated with cardiomyopathy-related genetic variants, highlighting the atrial myocardium as an important therapeutic target and understanding atrial-specific regulatory mechanisms provides new insights into therapeutic strategies for atrial diseases.

## Introduction

The heart is a vital organ responsible for maintaining sufficient perfusion of the body’s tissues. It achieves this through the coordinated function of its four chambers: the left and right atria (LA and RA, respectively) and the left and right ventricles (LV and RV, respectively). Each chamber has distinct structural and functional properties tailored to its role in circulation. The atria are thin-walled chambers that primarily function as reservoirs, passively filling the ventricles through elastic recoil. Active atrial contraction occurs only at the end of ventricular filling and contributes approximately ~20% of the total ventricular filling volume(1). In contrast, the ventricles are thick-walled, muscular chambers designed for forceful contractions that propel blood through the pulmonary and systemic circulations. In large mammals, including pigs and humans, atrial and ventricular myocardium differ not only in structure and function but also at the molecular level. Ventricular muscle predominantly expresses β-myosin heavy chain, which has a slower ATPase activity and supports slow, sustained, and forceful contractions (2). In contrast, atrial muscle contains a higher proportion of α-myosin heavy chain, which has a faster ATPase activity and favors rapid, burst-like contractions(2). Other sarcomeric thick filament proteins, including myosin light chains(3), titin(4), myosin-binding protein C(5), also exhibit chamber-specific isoforms that may contribute to differences in mechanical properties. Together, these chamber-specific differences in thick filament protein composition raise the possibility that LA and LV also differ in the structural regulation of thick filament activation. Because thick filament activation mechanisms are known to differ between fast- and slow-twitch skeletal muscle (6, 7), we hypothesized that analogous differences exist between fast atrial and slow ventricular porcine myocardium.

To test this hypothesis, we used small-angle X-ray diffraction to investigate thick filament structural and regulatory differences between porcine LA and LV across increasing calcium concentrations. We show that, at diastolic calcium levels, the thick filament in LA adopts a more ON state compared with those in LV. We then examined thick filament structural similarities and differences between LA and LV during calcium-activated contraction. To further isolate calcium-dependent regulation from the effects of active force generation, we employed the troponin inhibitor MYK-7660 to suppress contraction and selectively probe thick filament activation (8). Under these conditions, in contrast to ventricular myocardium, atrial myocardium does not exhibit calcium-dependent thick filament activation. This finding may help reconcile previously conflicting observations from ventricular studies in porcine (8, 9) and rodent (10) ventricular myocardium having slow and fast isoforms of myosin respectively. Collectively, these findings demonstrate that atrial myocardium in large mammals adopts a distinct structural configuration and regulatory mechanism optimized for rapid, burst-like contractions, whereas ventricular myocardium is tuned for sustained, energy-efficient force generation.

Historically, research and therapeutic development have predominantly focused on the ventricular muscle due to its central role in driving cardiac output, whereas atrial structure and function have received comparatively less attention. However, atrial dysfunction is increasingly recognized as a contributor to cardiac disease, including arrhythmias and heart failure. A substantial proportion of these atrial diseases carry cardiomyopathy-associated variants (11–13), making the atrial myocardium a potential target for emerging therapeutic strategies (14). By defining these chamber-specific regulatory mechanisms, this work provides a basis for understanding how different cardiac chambers adapt to their physiological roles, how atrial dysfunction may arise in disease, and how these insights may guide the development of targeted therapeutic strategies.

## Materials and Methods

### Ca^2+^ solutions

Ca^2+^ solutions were prepared as previously described (15). Free Ca^2+^ concentrations ranged from 0.01 μM (resting) to 46.8 μM (maximal activation; 5.95 mM Na_2_ATP, 6.20 mM MgCl_2_, 10 mM Ca-EGTA, 100 mM BES, 10 mM creatine phosphate, 29.98 mM K-propionate, protease inhibitor (Sigma-Aldrich, St. Louis, MO), and 1 mM DTT; pH 7.0), with the latter corresponding to approximately pCa 4.3. All solutions contained 3% Dextran T-500.

### Cardiac sample preparation

Porcine LA and LV were isolated from and prepared as previously described (16). Samples were permeabilized with 1% Triton X-100 for 4 h at 4 °C. Preparations were mounted between a force transducer (model 403A, Aurora Scientific Inc., Ontario, Canada) and a high-speed servomotor (model 312C, Aurora Scientific Inc.) for force measurement and length control. Steady-state isometric Ca^2+^-dependent force was measured by sequential exposure of the samples to a series of Ca^2+^ solutions described above at ~30 °C. Sarcomere length was set to 2.3 μm in pCa 8 using HeNe laser diffraction. Samples were then incubated in pCa 8 containing 3% dextran T-500 for 1 h before experiments.

### Muscle mechanics

Relative force–pCa relationships were fitted in GraphPad Prism using a two-parameter Hill equation, whereas absolute force–pCa relationships were fitted using a four-parameter Hill equation. For relative force, data were fitted as:

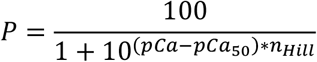

where *P* is relative force (%), *pCa*_*50*_ is the pCa producing 50% of maximal force, and *n*_*Hill*_ is the Hill coefficient. For absolute force (mN/mm^2^), data were fitted as:

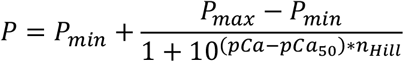

Where *P* is force, and *P*_max_ and *P*_min_ represent the upper and lower plateaus of the force–pCa relationship, respectively.

The kinetics of tension redevelopment (*k*_TR_) were measured after isometric force had reached steady state at each level of Ca^2+^ activation. *k*_TR_ was determined by rapidly shortening CMPs to 20% of their initial length (L_0_), followed by a rapid re-stretch to 25% beyond the original length and subsequent shortening back to L_0_. *k*_TR_ was calculated from individual recordings as previously described (17). Sinusoidal stiffness (SS) was measured after isometric force had reached steady state in each pCa solution by recording changes in sarcomere length and the corresponding changes in force. CMPs were subjected to small-amplitude length oscillations of 0.2% peak-to-peak of the initial length at a frequency of 100 Hz and a sampling rate of 1 kHz (18). SS data were analyzed in RStudio using Fast Fourier transform, and values were reported in MPa. SS-pCa relationships were fitted using a four-parameter sigmoidal Hill equation, as described above. All experiments were performed at 30 °C.

### Mathematical modeling

The relationship between steady-state isometric force and *k*_TR_ was modeled using a three-state model of muscle contraction (17). The coupled differential equations describing the model were solved in MATLAB for: *f, g, k*_ON_, and *k*_OFF_. *f, g*, and *k*_OFF_ were iteratively varied from their initial estimates to identify the best-fit values using the Simplex algorithm, while *k*_ON_ was held constant based on values reported by Pinto et al (19). Best-fit parameter estimates were determined using a least-squares approach. Specifically, the sum of squared deviations (SSD) was minimized by accounting for both the difference between predicted and measured force– *k*_TR_ relationships and the difference between predicted and measured pCa_50_ values for isometric force. Individual SSD components were weighted to prioritize maximum *k*_TR_, pCa_50_, and maximum force normalized to LV.

### Characterization of MYK7660 on LA

The dose-dependent maximally activated force (pCa4.3) data are collected in the presence of increasing MYK-7660 concentrations (0 μM, 1 μM, 10 μM, 25 μM, 50 μM, 100 μM and 180 μM in both pCa 8 and pCa 4.3 solutions) on skinned porcine LA (Fig S1B). The tissues were incubated in pCa 8 solution containing MYK-7660 for 10 min before activation in pCa 4.3 solution. The tissues were then relaxed in pCa 8 solution containing the next concentration of MYK-7660 for 10 min prior to subsequent activation at pCa 4.3. Forces are normalized against the force generated with no inhibitor.

### Muscle preparation for X-ray diffraction

Wild-type porcine hearts were purchased from Exemplar Genetics. The hearts were cut into smaller pieces and frozen by liquid nitrogen and stored in −80°C freezer as described previously (16). Frozen left ventricular (LV) and left atrial (LA) myocardium were thawed in skinning solution (2.25 mM Na_2_ATP, 3.56 mM MgCl_2_, 7 mM EGTA, 15 mM sodium phosphocreatine, 91.2 mM Potassium Propionate, 20 mM Imidazole, 0.165 mM CaCl_2_, 15 mM 2,3-Butanedione 2-monoxime (BDM),1% Triton X-100 and protease inhibitor cocktail) for ~30 minutes at room temperature before splitting them into smaller fiber bundles for further skinning either for 2-3 hours at room temperature or overnight at 4°C on a rocker. Once the skinning is complete, the muscles were washed with fresh low calcium (pCa8.0) cardiac relaxing solution (2.25 mM Na_2_ATP, 3.56 mM MgCl_2_, 7 mM EGTA, 15 mM sodium phosphocreatine, 91.2 mM Potassium Propionate, 20 mM Imidazole, 0.165 mM CaCl_2_ and protease inhibitor cocktail) three times, 10 minutes each. The muscle bundles were further dissected into well-aligned, unified bundles (~300μm in diameter) and clipped with aluminum T-clips and stored in cold relaxing containing 3% dextran on ice for use in experiments on the same day.

### X-ray Diffraction

X-ray diffraction experiments were performed at the BioCAT beamline 18ID at the Advanced Photon Source, Argonne National Laboratory (20) and at the HP BioSAXS & BioSAXS 7A beamline at the Cornell High Energy Synchrotron Source (CHESS). The X-ray beam energy was set to 12 keV (0.1033 nm wavelength). The specimen-to-detector distance was ~3m at BioCAT and ~1.7m at HP BioSAXS & BioSAXS. The preparation was then attached at one end to a hook on a force transducer (Model 402B Aurora Scientific Inc., Aurora, ON, Canada) and the other end to a static hook. The muscles were stretched to a sarcomere length of 2.3 µm by helium-neon laser (633 nm) diffraction and the solution temperature was maintained between 28°C to 30°C by a heat exchanger attached underneath the chamber. The X-ray patterns are collected sequentially from both LV and LA myocardium (Fig 3) at six increasing Ca^2+^ concentrations (pCa 8, pCa 6.8, pCa 6.2, pCa 5.8, pCa 5.4 and pCa 4.5) using a MarCCD 165 detector (Rayonix Inc., Evanston IL) with a 1 s exposure time at BioCAT and on an Eiger 4M detector (Dectris, Switzerland) with a 2 s exposure time at HP BioSAXS & BioSAXS. X-ray diffraction patterns were also collected from LA myocardium at the same increasing Ca^2+^ concentrations in the presence of 180 μM of MYK-7660 to prevent active contraction as Ca^2+^ concentrations increase. No noticeable force was observed throughout this experiment. The muscle samples were oscillated horizontally at a speed of 1–2 mm/s during data collection and following each exposure, the sample was shifted vertically to reduce radiation damage.

### X-ray data analysis

The data are analyzed using MuscleX software package developed at BioCAT (21). The equatorial reflections are measured by the “Equator” routine in MuscleX as previously described (22). The X-ray patterns were then quadrants folded and background subtracted to improve the signal-to-noise ratio using the “Quadrant Folding” routine in MuscleX. The meridional and layer line reflections were analyzed by the “Projection Traces” routine in MuscleX as described previously (23). The measured intensities of X-ray reflections are normalized to the sixth-order actin-based layer line intensities whose intensity did not change significantly under various condition (8, 24). Two to three patterns are collected under each condition, and reflection spacings and intensities extracted from these patterns are averaged.

### Statistics

Statistical analyses are performed using GraphPad Prism 11.0.1 (Graphpad Software). The results are given as mean ± SEM. Statistical significance in mechanics experiments was assessed by unpaired Student’s t-test. The X-ray signal values versus pCa curves are fit to a four-parameter modified Hill equation (Y=Bottom + (Top-Bottom)/(1+10^((LogEC50-X)*HillSlope) in Fig 3 and Fig4. Difference between groups at the same pCa were compared using a mixed-effects model with restricted maximum likelihood (REML) estimation in place of grouped two-way ANOVA due to missing values in the dataset. Post hoc multiple-comparisons testing was performed using Sidak’s correction. The inset bar graphs in Fig 3A and Fig 3F were performed using unpaired t test with Welch’s correction. Symbols on figures: *: p<0.05, **: p<0.01, ***: p<0.001, ****: p<0.0001.

## Results

### Porcine LA and LV exhibit distinct force generating capacity and cross-bridge cycling kinetics

When steady-state isometric force was compared between porcine LV and LA, LA developed significantly lower force than LV across most activating calcium concentrations (Fig.1 A). This reduction was evident beginning at pCa 6.1 and extending through maximal calcium activation (pCa 4.3). At maximal calcium activation, LA generated 69% of the force produced by LV (61.04 ± 6.81 mN/mm^2^ vs 88.47 ± 5.10, *p* < 0.01, Table S1). At lower calcium concentrations, including pCa 8.0, 7.0, and 6.5, LA force tended to be lower than LV, although these differences did not reach statistical significance (Fig.1 A). Thus, the main difference between chambers was observed at systolic calcium levels and above, where LA generated less force than LV. Despite these differences in steady-state isometric force generation, myofilament calcium sensitivity (pCa_50_, LA = 5.89 ± 0.02 vs LV = 5.95 ± 0.04, *p* > 0.05; Table S1, Fig.1 B and C) and the cooperativity of thin filament activation (*n*_Hill_, LA = 2.32 ± 0.20 vs LV = 2.39 ± 0.25, *p* > 0.05; Table S1, Fig.1 B and D) were not significantly different between LV and LA.

**Figure 1.**
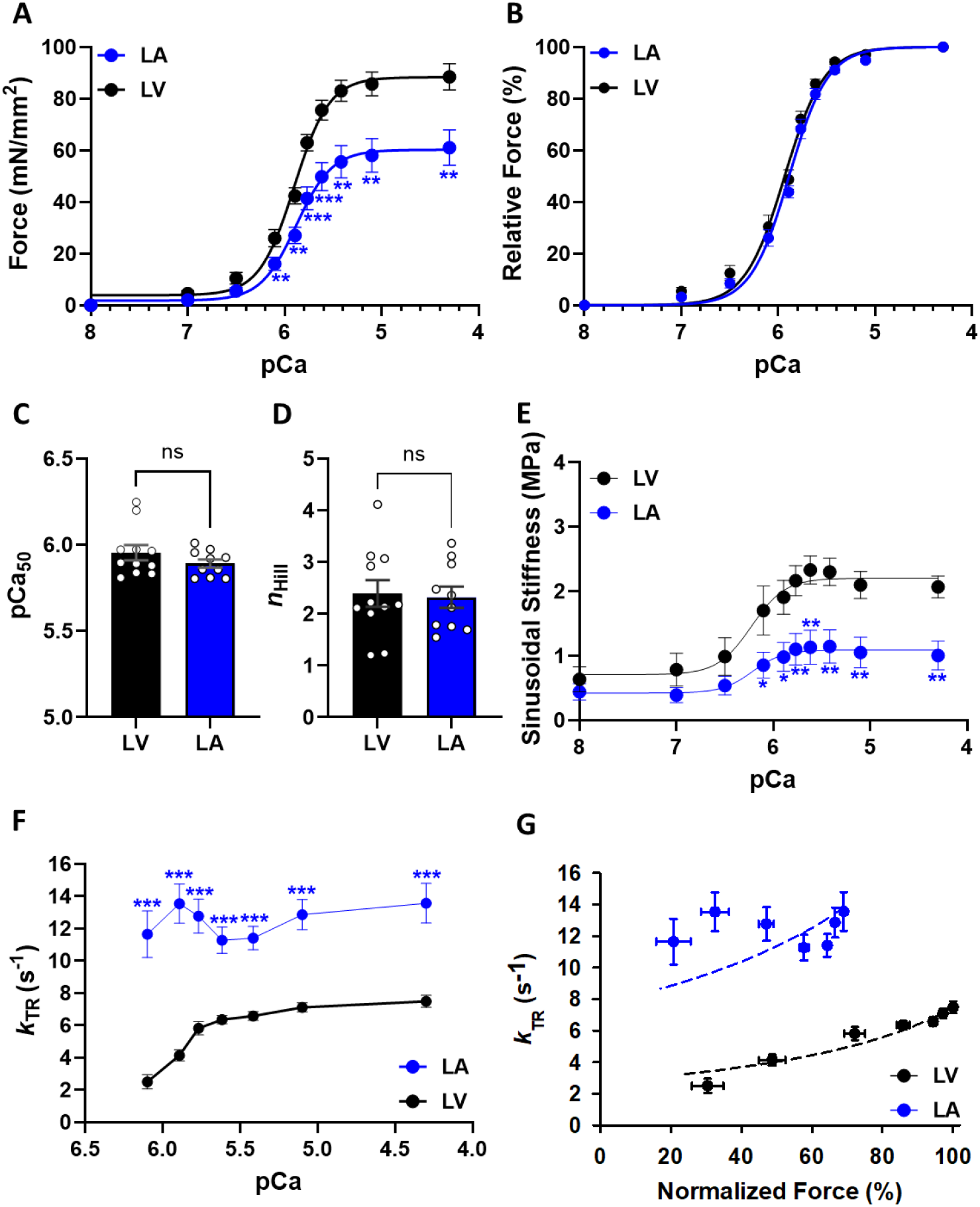
Contractile properties of porcine left atrial (LA) and ventricular (LV) myocardium. **A)** pCa-force curves at sarcomere length (SL) 2.3 µm. Force levels are normalized to the cross-sectional area of each cardiac muscle preparations. **B)** pCa-relative force plot at SL 2.3 µm: The force levels are relative to the maximum steady-state isometric force generated by each preparation. **C)** Myofilament calcium sensitivity (pCa_50_): the amount of calcium necessary to reach 50% of the force. **D)** Cooperativity of thin filament activation. **E)** pCa-sinusoidal stiffness (SS) curves at SL 2.3 µm. Megapascal is the unit for SS. **F)** pCa-kinetics of tension redevelopment (*k*_TR_) curves at SL 2.3 µm. **G)** Normalized force-*k*_TR_ curves at SL 2.3 µm. LV force was normalized by the maximum force generated by each LV preparation. LA force was normalized by the average maximum steady-state isometric force generated by LV. Dash lines represent the solutions to the 3-state mathematical model using the best fit parameters (*f, g, k*_OFF_) and *k*_ON_ listed in Table S2.

We next assessed sinusoidal stiffness (SS), which provides an estimate of the overall cross-bridge population. Similar to the steady-state isometric force measurements, SS was significantly lower in LA than in LV (Fig.1 E) at the same calcium concentrations where force was reduced (Fig.1 A and Table S1). At lower calcium concentrations, LA stiffness also tended to be lower than LV stiffness, although these differences were not statistically significant (Fig.1 E). Kinetics of tension redevelopment (*k*_TR_), were markedly faster in LA than in LV across all calcium concentrations tested (Fig.1 F and Table S1). LV exhibited a clear calcium (Fig.1 F) or force (Fig.1 G) dependence of *k*_TR_, while LA showed much less calcium dependence. Last, we used mathematical modeling to estimate the apparent cross-bridge attachment rate (*f*), detachment rate (*g*), and the rate of calcium dissociation from the thin filament (*k*_OFF_) (19). The model predicted that LA has both higher *f* and higher *g* compared with LV (LA, *f* = 6.16 and *g* = 7.81; LV, *f* = 4.87 and *g* = 2.77; Table S2). Notably, in LA, *g* exceeded *f*, whereas in LV, *f* was greater than In addition, *k*_OFF_ was higher in LV than in LA (LA, *k*_OFF_ = 456; LV, *k*_OFF_ = 581; Table S2).

### Left atrial and ventricular thick filament structural arrangements at different levels of calcium

Under resting conditions, a large fraction of myosin heads adopts a quasi-helical, ordered OFF state, in which the heads are closely aligned along the surface of the thick filament backbone. This highly ordered organization gives rise to the characteristic strong X-ray diffraction signals, particularly the myosin-based layer lines (MLLs), observed in resting muscle (25, 26). In contrast, perturbations such as muscle activation (26, 27), small-molecule treatment (13, 28, 29), physiological regulation (30), or pathological remodeling (31) can disrupt this order, shifting myosin heads into a more disordered ON state in which they extend away from the thick filament backbone (26). Under low-calcium conditions, porcine LV exhibits a characteristic 2D X-ray diffraction pattern with well-defined equatorial, meridional, and layer-line reflections (Fig. 2A, right panel) (29, 32). LA also displays full 2D diffraction patterns (Fig. 2B, right panel); however, the MLLs are qualitatively weaker than those observed in ventricular myocardium under low-calcium conditions, whereas the intensity of the sixth-order actin layer line (ALL6) is comparable. To the best of our knowledge, this study provides the first report of X-ray diffraction patterns obtained from atrial myocardium. Upon an increase of calcium concentration, the intensity of the MLLs is markedly reduced at pCa 4.5, while the ALL6 remains relatively constant in both muscle types (Fig 2, left panel). The equatorial intensity ratio (I_1,1_/I_1,0_) indicates myosin head proximity to actin in relaxed muscle and quantitatively reflects the number of force-generating cross-bridge upon activation(26). At resting (pCa 8), I_1,1_/I_1,0_ was significantly lower in LV than in LA (0.23 ± 0.02 vs. 0.32 ± 0.02, respectively, *p* = 0.005, inset of Fig 3A). This finding indicates that, under relaxing conditions, atrial myosin heads are positioned farther from the thick filament backbone and/or closer to actin compared to ventricular myosin heads. Increasing calcium concentration produced a sigmoidal rise in I_1,1_/I_1,0_ for both chambers. In LV, the I_1,1_/I_1,0_ pCa_50_ was 5.76 with a maximal I_1,1_/I_1,0_ of 1.05 ± 0.3 at pCa 4.5 (Fig. 3A). In LA, I_1,1_/I_1,0_ pCa_50_ was 5.45, and the maximal I_1,1_/I_1,0_ reached 2.99 ± 0.8 at pCa 4.5 (Fig. 3A). At pCa 8, the 1,0 lattice spacing (d_1,0_) was significantly larger in LA than in LV (37.2 ± 0.19 nm vs. 36.4 ± 0.34 nm, respectively; *p* = 0.026; Fig. 3B). With increasing calcium concentration, d_1,0_ increased in a sigmoidal fashion, whereas atrial d_1,0_ remained largely unchanged.

**Figure 2.**
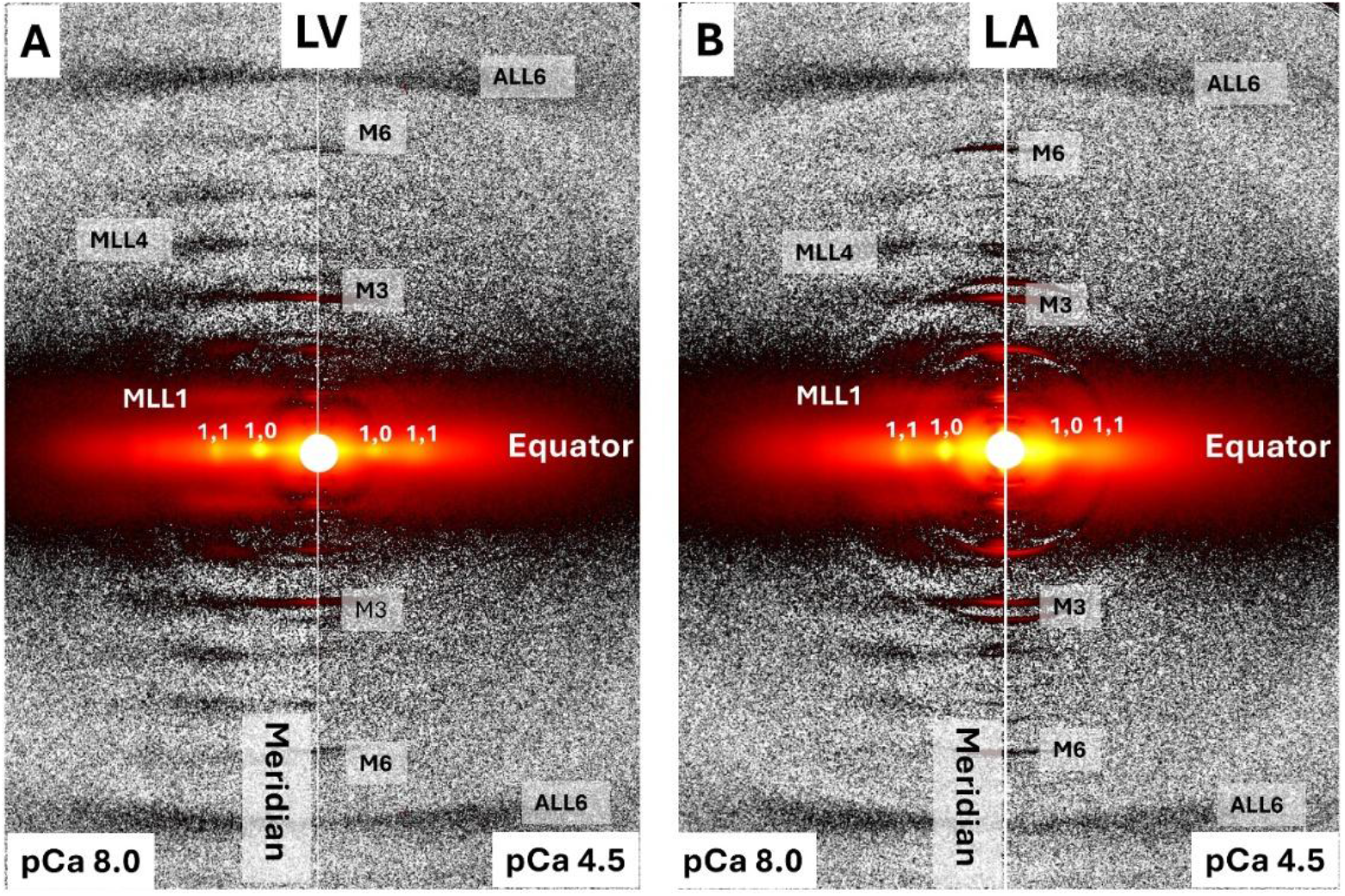
Representative X-ray diffraction patterns from permeabilized porcine left ventricular and atrial myocardium. X-ray diffraction patterns from permeabilized porcine **A)** left ventricular and **B)** atrial myocardium under low (pCa 8.0) and high (pCa 4.5) calcium conditions.

**Figure 3.**
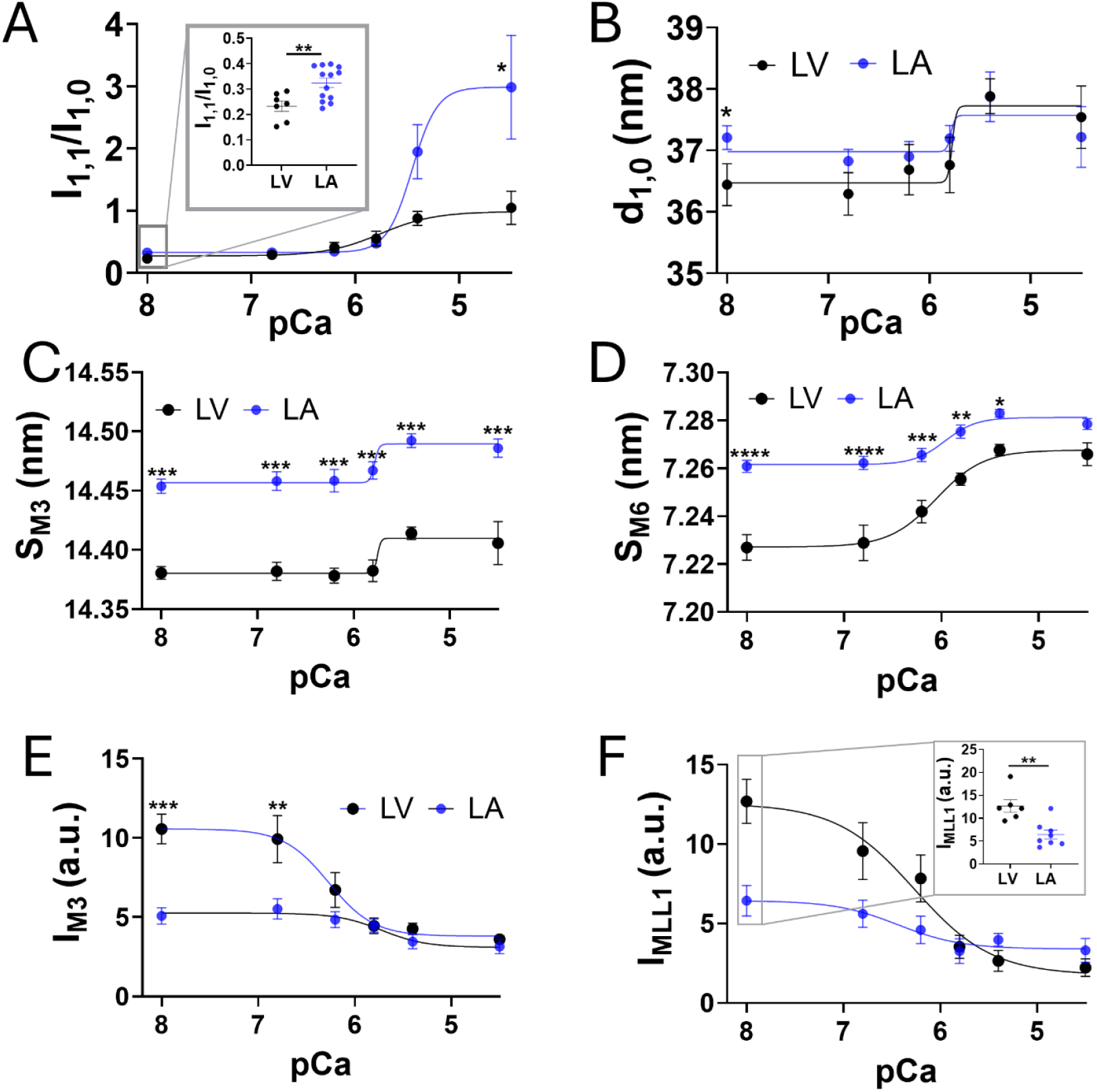
Thick filament structural changes at increasing levels of calcium from left ventricular and atrial myocardium. Equatorial intensity ratio (I_1,1_/I_1,0_, **A**) and 1,0 lattice spacing (d_1,0_, **B**) at different calcium concentrations from left ventricular (LV, black) and atrial (LA, blue) myocardium. The spacing of the third-order (S_M3_, **C**) and sixth-order (S_M6_, **D**) of myosin-based meridional reflection in different Calcium concentrations in LV and LA. The intensity of the third-order myosin-based meridional reflection (I_M3_, **E**) and first-order myosin-based layer-line (I_MLL1_, **F**) of LV and LA in different Calcium concentrations.

The spacing of the third-order myosin-based meridional reflection (S_M3_), which reflects the axial periodicity of myosin heads, was significantly greater in LA than in LV across the full range of calcium concentrations (Fig 3C). This finding indicates distinct myosin head configurations in atrial muscle under both relaxing and activation conditions compared with ventricular myosin heads. Similarly, the spacing of the sixth-order myosin-based meridional reflection (S_M6_), which is customarily attributed to periodicities within the thick filament backbone (27, 33), was consistently larger in LA than in LV, suggesting intrinsic differences in backbone conformation between the two chambers (Fig 3D). The intensity of the third-order myosin-based meridional reflection (I_M3_) and the first-order myosin-based layer line (I_MLL1_) correlate with the ordering of myosin heads (25, 34). Under relaxing condition, these parameters were about two-fold higher in LV than in LA (I_M3_: 10.57 ± 0.94 in LV vs. 5.09 ± 0.51 in LA (Fig 3E), *p* = 0.0002; I_MLL1_: 12.69 ± 1.39 in LV vs. 6.45 ± 0.97 in LA (Fig 3F), *p* = 0.002), indicating that myosin heads in LA adopt a more disordered configuration compared with LV. Both I_M3_ and I_MLL1_ decreased progressively with increasing calcium concentration, and the differences between LV and LA became progressively attenuated.

### Direct calcium-mediated activation of the thick filament in porcine myocardium

Previous studies have shown that porcine ventricular myocardium exhibits calcium-dependent activation of the thick filament that occurs independently of cross-bridge–mediated force generation (8). This mechanism was examined in isolation by pharmacologically inhibiting thin filament–based force production using the troponin inhibitor MYK-7660, thereby uncoupling calcium-dependent thick filament activation from calcium-dependent force development (8). Here, we used the same troponin inhibitor to assess calcium-dependent thick filament activation in porcine LA and to compare these responses with previous findings in porcine LV, where direct calcium-dependent thick filament activation has been observed (8). In LV, the change in I_1,1_/I_1,0_ (_Δ_I_1,1_/I_1,0_) as a function of pCa showed a similar sigmoidal profile in both control and inhibitor-treated groups (Fig 4A). Under control conditions, Δ I_1,1_/I_1,0_ reached 0.64 ± 0.08, whereas this response was blunted to 0.27 ± 0.02 in the inhibitor-treated group. Thus, in the absence of thin filament-mediated force production, the calcium-dependent chance in Δ I_1,1_/I_1,0_ was preserved at approximately 47% of the control response in LV. In contrast, LA showed a markedly different response. Although Δ I_1,1_/I_1,0_ reached 2.66 ± 0.85 under control conditions, it increased only by 0.075 ± 0.034 in the presence of the troponin inhibitor, corresponding to merely 3% of the LA control response (Fig 4B). Similarly, I_M3_ progressively decreased with increasing calcium concentration in control LA (Fig 4D), resembling the pattern observed in LV (Fig 4C). However, in the presence of the inhibitor, I_M3_ remained unchanged in LA across calcium concentrations (Fig 4D). Increases in S_M6_, which reports the periodicity of the thick filament backbone, may be attributed to both the strain imposed on the thick filament during active contraction and to the OFF-to-ON transitions of myosin heads within the thick filament (8, 24, 35). In LV, S_M6_ increased by 0.73 ± 0.06% at full activation (pCa 4.5) under control conditions and by 0.36 ± 0.02% in the inhibitor-treated group, in which active force was abolished (Fig 4E). These findings indicate that approximately half of the S_M6_ increase in LV can be attributed to the OFF-to-ON transition of the thick filament. In LA, S_M6_ increased by 0.35 ± 0.04% at pCa 4.5 under control conditions, whereas S_M6_ showed no significant calcium-dependent increase in the inhibitor-treated group (Fig 4F). Together, these results show that, unlike LV, LA does not exhibit a calcium-dependent OFF-to-ON transition of the thick filament when thin filament-mediated force-production is inhibited.

**Figure 4.**
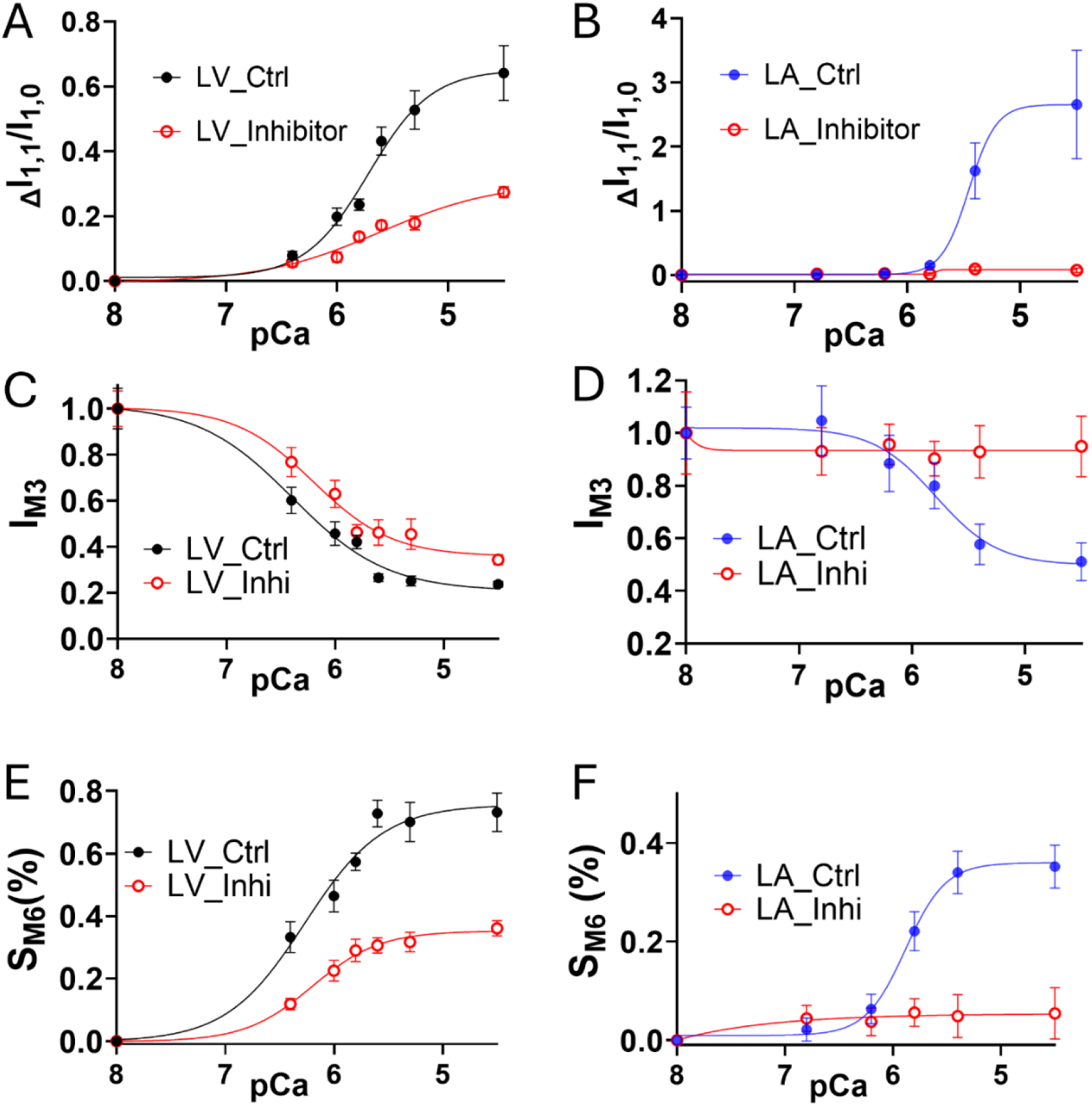
Calcium-dependent thick filament structural changes in left ventricular (left panels, replotted from(8)) and atrial (right panels) myocardium in the absence of active contraction. Change of equatorial intensity ratio (ΔI_1,1_/I_1,0_) in different calcium concentrations from LV (**A**) and LA (**B**) myocardium. Normalized I_M3_ in different calcium concentrations from LV (**C**) and LA (**D**) myocardium. Change of S_M6_ in different calcium concentrations from LV (**E**) and LA (**F**) myocardium.

## Discussion

### Distinct structural configurations of sarcomeric proteins in porcine LA and LV under resting conditions

Muscle contraction is tightly regulated, traditionally thought to primarily consist of thin filament-based mechanisms, but are now understood to include thick filament–based regulatory mechanism(s) that operate in parallel (36, 37). Myosin heads exist in two main structural states: an ordered OFF state, which serves as a reserve pool, and a disordered ON state, which is capable of interacting with actin and contributing to contraction (25, 37). The equilibrium between the OFF/ON state of the myosin heads in the resting state plays an essential role in determining the magnitude of subsequent calcium-activated force (24). The signatures from our X-ray diffraction pattern under low calcium concentration (pCa 8) showed that myosin heads in porcine LA adopts a different configuration compared to LV under the same conditions. Specifically, I_1,1_/I_1,0_ in LA was larger than in LV, indicating that atrial myosin heads are positioned farther from the thick filament backbone and exhibit greater association with the actin filament than those in the ventricles. Additionally, I_M3_ and I_MLL1_, indicators of the degree of ordering of myosin heads, were about half the values in LA and LV, indicating that myosin heads in LA adopt a more disordered configuration during diastole compared with those in the LV. Moreover, S_M3_ and S_M6_, whose increase is strongly associated with myosin heads transitioning to the ON state, were significantly greater in LA. Altogether, our X-ray data indicate that myosin filaments in LA adopt a more ON-like and structurally distinct configuration compared to those in LV, a feature that may be evolutionarily advantageous in meeting the rapid contractile demands of the atrium. The molecular basis of this phenomenon may lie in the observation that the abundance of cardiac myosin-binding protein C (cMyBP-C) in human atrial myocardium is approximately half that found in ventricular myocardium (38). In the atrial sarcomere, roughly half of the canonical cMyBP-C binding sites appear to be occupied by myosin-binding protein H-like protein. cMyBP-C has been extensively shown to stabilize the OFF state of myosin (39, 40). Therefore, the reduced level of cMyBP-C in atrial myocardium could provide a molecular explanation for why a greater proportion of myosin heads adopt the ON configuration compared with those in ventricular myocardium.

### Distinct structural and functional properties of porcine LA and LV under contracting conditions

In our study, LV developed significantly greater force than the LA across all calcium concentrations (Fig 1A). The lack of difference in pCa50 and n_Hill_ indicate that the reduced force in LA is not explained by a major shift in calcium sensitivity or thin filament cooperative activation. Sinusoidal stiffness (SS) was significantly lower in LA than in LV, suggesting that the reduced force in the LA may be associated with a smaller population of cross-bridges. Additional factors, including the lower myofilament density in atrial tissue compared with ventricular tissue (2), may also contribute to the reduced contractile force in the LA. Previous studies comparing LA and LV have reported mixed findings. van der Velden et al (41), and Nakanishi et al (42) reported higher tension development in LV compared with LA in human and porcine tissue, respectively, whereas Narolska et al. (43) and Piroddi et al (44) observed the opposite result in human myocardium. The exact reasons for these discrepancies remain unclear. However, differences in experimental conditions may contribute to these discrepancies, including variations in temperature, sarcomere length, and the use of dextran to restore myofilament lattice spacing altered during permeabilization to near *in vivo* values. Our mechanical data also showed that calcium sensitivity (pCa50) was similar between the LV and LA, consistent with the studies mentioned above. The *k*_TR_ was markedly higher in the LA than in the LV across all pCa levels, suggesting intrinsic differences in cross-bridge cycling kinetics between fast α-myosin in the LA and slow β-myosin in the LV. Interestingly, *k*_TR_ appeared largely insensitive to calcium concentration in the LA, whereas it increased with rising calcium in the LV, suggesting a unique calcium-dependent modulation of cross-bridge cycling kinetics that is specific to slow β-myosin.

Structurally, one distinct difference in X-ray signatures during contraction was the markedly greater increase in the I_1,1_/I_1,0_ in the LA compared with the LV at high calcium concentrations (pCa 5.4 and pCa 4.5, (Fig 3A)). The maximal I_1,1_/I_1,0_ was 2.99 ± 0.8 in the LA, approximately threefold higher than that observed in the LV. Such high I_1,1_/I_1,0_ values are more commonly observed in fully contracting skeletal muscle patterns (22, 23, 45). The higher S_M3_ (Fig. 3C) and S_M6_ (Fig. 3D) values in the LA were maintained as calcium concentration increased, indicating that the distinct myosin filament configuration in the LA persisted during contraction. In contrast, the differences in I_M3_ and I_MLL1_ diminished with increasing calcium concentration, suggesting that the additional myosin heads in the OFF state in the LV were progressively recruited, reaching a level similar to that in the LA at high calcium concentrations. Collectively, these findings demonstrate distinct functional properties of the LA and LV in force generation, cross-bridge recruitment, and cross-bridge cycling kinetics, as well as distinct structural properties in myosin heads configuration during contraction.

### Porcine LA showed no calcium-dependent direct thick filament activation

The mechanisms by which myosin heads transition from the OFF to the ON state, i.e. thick filament–based regulation, have not yet been fully elucidated. The most widely accepted explanation is the mechanosensing model where the thick filament functions as a strain sensor that responds to tension generated by a small number of constitutively ON cross-bridges. This mechanical strain promotes the release of additional myosin heads from the OFF state into the ON state, thereby amplifying force production during contraction (27, 46, 47). The mechanosensing model is primarily derived from fast skeletal muscle (27) and rodent cardiac muscle (46, 47). Subsequent studies in slow skeletal muscle, however, showed that the characteristic X-ray diffraction signatures of the mechanosensing model are attenuated, suggesting that additional or alternative mechanisms of thick filament regulation may be required (6). It has been shown that calcium may directly activate the thick filament in porcine left ventricular myocardium, and the extent of the OFF-to-ON transition remains comparable even when mechanosensing is abolished by inhibiting thin filament activation and thereby preventing active force generation (8, 9). Kalakoutis et al.(10), using the same methodology, reported contrasting results from rat ventricular myocardium, showing that the thick filament does not respond to calcium in the absence of calcium binding to the thin filament. Here, we examined the direct effects of calcium on thick filament activation in porcine LA which differs from porcine LV but, like rat ventricular myocardium, predominantly expresses the α-myosin heavy chain isoform. Surprisingly, porcine LA showed no evidence of direct calcium-dependent thick filament activation and instead behaved similarly to rat ventricular myocardium. In the presence of MYK-7660, all signature structural changes associated with the OFF-to-ON transition of the thick filament (i.e., I_1,1_/I_1,0_, I_M3_, and S_M6_) were absent as calcium concentration increased. These findings suggest that direct calcium-mediated activation of the thick filament is not a universal property of cardiac muscle but instead depends on the molecular composition of the thick filament, potentially reflecting adaptation to specific mechanical demands. One possible determinant is the myosin isoform composition: porcine LA and rat ventricle predominantly express the α-myosin heavy chain isoform, whereas porcine ventricular myocardium predominantly expresses the β-myosin heavy chain isoform. Together, these observations suggest that myosin isoform composition may be a primary determinant, along with other contributing factors, of the structural and regulatory responsiveness of thick filament activation mechanisms. This distinction may help reconcile the seemingly contradictory findings between porcine (8, 9) and rat ventricular (10) studies.

### Thick filament regulation has evolved to match the specific functional demands of each muscle type

One might ask why atrial myocardium exhibits different thick filament regulation than ventricular myocardium in the porcine heart. However, given the distinct function, morphology, and molecular composition of the atrium and ventricle, the more appropriate question may be why they should be the same. Mechanosensing is a mechanism that optimizes both speed and force, enabling rapid muscle activation suited for fast, burst-like movements (46). The atrium contracts only at the end of ventricular filling, where the primary goal is to maximize ventricular preload (5). Thus, a greater proportion of myosin heads in the ON state would be advantageous, enhancing force generation and optimizing atrial function. In this context, speed and force are prioritized over precision. In contrast, the ventricle’s primary role is to meet the body’s varying demands for oxygen and nutrients, where precise control of force generation is essential. Direct calcium-dependent activation of the thick filament may support this fine-tuned regulation in large mammalian hearts, which typically operate at only ~10% of their maximal functional capacity (48). In addition, maintaining a large molecular reserve—where a substantial fraction of myosin heads remains in the OFF state—may be advantageous, providing recruitment capacity when increased contractile capacity is required. In comparison, this mechanism may be less critical in rodent hearts, which operate at a much higher baseline workload (~80% of maximal capacity) and at heart rates 5–10 times higher than those of large mammalian hearts (49). Together, these differences underscore that thick filament regulation is not the same in all tissues and in all organisms but may be shaped by the specific functional demands of each muscle type.

## Author contributions

W.M, T.C.I, and S.N designed the experiments; H.F, S.Y, M.L-V, E.O.O, H.G, M.M and W.M performed the experiments; L.Q, M.L-V, M.G, and C.G analyzed the data; M.L-V and W.M wrote the initial draft. All authors approved the final version of the manuscript.

## Acknowledgments

This project is supported by National Heart Lung and Blood Institute Grant (R01HL171657 and 1R01HL172492). This research used resources of the Advanced Photon Source, a U.S. Department of Energy (DOE) Office of Science User Facility operated for the DOE Office of Science by Argonne National Laboratory under Contract No. DE-AC02-06CH11357. BioCAT is supported by grant P30 GM138395 from the National Institute of General Medical Sciences of the National Institutes of Health. The content is solely the authors’ responsibility and does not necessarily reflect the official views of the National Institute of General Medical Sciences or the National Institutes of Health

## Data Availability

The dataset generated or analyzed during this study is included in this article. The raw data are available from the corresponding authors (Weikang Ma) on reasonable request.

## References

1. R. E. Klabunde, Cardiovascular physiology concepts (Wolters Kluwer, Philadelphia, ed. Third edition., 2022), pp. xi, 265 pages.

2. J. Walklate et al., Alpha and beta myosin isoforms and human atrial and ventricular contraction. Cell Mol Life Sci 78, 7309–7337 (2021).

3. Z. R. Gregorich et al., Distinct sequences and post-translational modifications in cardiac atrial and ventricular myosin light chains revealed by top-down mass spectrometry. Journal of molecular and cellular cardiology 107, 13–21 (2017).

4. O. Cazorla et al., Differential expression of cardiac titin isoforms and modulation of cellular stiffness. Circulation research 86, 59–67 (2000).

5. H. V. Burnham, H. E. Cizauskas, D. Y. Barefield, Fine tuning contractility: atrial sarcomere function in health and disease. American journal of physiology. Heart and circulatory physiology 326, H568–H583 (2024).

6. H. M. Gong, W. Ma, M. Regnier, T. C. Irving, Thick filament activation is different in fast-and slow-twitch skeletal muscle. The Journal of physiology 600, 5247–5266 (2022).

7. C. Hill et al., Distinct distributions of myosin motor conformations during contraction of slow and fast skeletal muscle. The Journal of physiology 10.1113/JP290232 (2026).

8. W. Ma, S. Nag, H. Gong, L. Qi, T. Irving, Cardiac myosin filaments are directly regulated by calcium. J Gen Physiol 154 e202213213 (2022).

9. S. Mohran et al., Calcium has a direct effect on thick filament activation in porcine myocardium. J Gen Physiol 156 (2024).

10. M. Kalakoutis et al., Calcium binding to troponin C is required for activation of the myosin-containing thick filaments in rat cardiac trabeculae. Journal of molecular and cellular cardiology 205, 129–138 (2025).

11. O. B. Vad et al., Rare and Common Genetic Variation Underlying Atrial Fibrillation Risk. JAMA Cardiol 9, 732–740 (2024).

12. M. S. Kandola et al., Population-Level Prevalence of Rare Variants Associated With Atrial Fibrillation and its Impact on Patient Outcomes. JACC Clin Electrophysiol 9, 1137–1146 (2023).

13. O. B. Vad et al., Prevalence of deleterious variants in cardiomyopathy genes in early-onset atrial fibrillation. Eur J Hum Genet 10.1038/s41431-026-02119-5 (2026).

14. P. Karakasis et al., Atrial Cardiomyopathy in Atrial Fibrillation: Mechanistic Pathways and Emerging Treatment Concepts. J Clin Med 14 (2025).

15. V. Jani et al., Right Ventricular Sarcomere Contractile Depression and the Role of Thick Filament Activation in Human Heart Failure With Pulmonary Hypertension. Circulation 147, 1919–1932 (2023).

16. W. Ma et al., The structural and functional integrities of porcine myocardium are mostly preserved by cryopreservation. J Gen Physiol 155 (2023).

17. D. Gonzalez-Martinez et al., Structural and functional impact of troponin C-mediated Ca(2+) sensitization on myofilament lattice spacing and cross-bridge mechanics in mouse cardiac muscle. Journal of molecular and cellular cardiology 123, 26–37 (2018).

18. M. A. Marques et al., Anomalous structural dynamics of minimally frustrated residues in cardiac troponin C triggers hypertrophic cardiomyopathy. Chem Sci 12, 7308–7323 (2021).

19. J. R. Pinto et al., Strong cross-bridges potentiate the Ca(2+) affinity changes produced by hypertrophic cardiomyopathy cardiac troponin C mutants in myofilaments: a fast kinetic approach. The Journal of biological chemistry 286, 1005–1013 (2011).

20. R. Fischetti et al., The BioCAT undulator beamline 18ID: a facility for biological non-crystalline diffraction and X-ray absorption spectroscopy at the Advanced Photon Source. J. Synchrotron Radiat. 11,, 399–405 (2004).

21. J. Jiratrakanvong et al., MuscleX: data analysis software for fiber diffraction patterns from muscle. Journal of synchrotron radiation 10.1107/S1600577524006167 (2024).

22. W. Ma, H. Gong, T. Irving, Myosin Head Configurations in Resting and Contracting Murine Skeletal Muscle. Int J Mol Sci 19 (2018).

23. W. Ma et al., Myosin dynamics during relaxation in mouse soleus muscle and modulation by 2’-deoxy-ATP. The Journal of physiology 598, 5165–5182 (2020).

24. W. Ma et al., Structural OFF/ON transitions of myosin in relaxed porcine myocardium predict calcium-activated force. Proceedings of the National Academy of Sciences of the United States of America 120, e2207615120 (2023).

25. N. A. Koubassova et al., Annotating the x-ray diffraction pattern of vertebrate striated muscle. Biophysical journal 10.1016/j.bpj.2025.09.019 (2025).

26. W. Ma, T. C. Irving, Small Angle X-ray Diffraction as a Tool for Structural Characterization of Muscle Disease. Int J Mol Sci 23 (2022).

27. M. Linari et al., Force generation by skeletal muscle is controlled by mechanosensing in myosin filaments. Nature 528, 276–279 (2015).

28. V. P. Jani et al., The structural OFF and ON states of myosin can be decoupled from the biochemical super-and disordered-relaxed states. PNAS Nexus 3, pgae039 (2024).

29. W. Ma et al., Myosin in autoinhibited off state(s), stabilized by mavacamten, can be recruited in response to inotropic interventions. Proceedings of the National Academy of Sciences of the United States of America 121, e2314914121 (2024).

30. W. Ma et al., The Super-Relaxed State and Length Dependent Activation in Porcine Myocardium. Circulation research 129, 617–630 (2021).

31. S. Steczina et al., Molecular mechanisms of altered contraction with the beta-myosin R403Q mutation in porcine ventricular muscle and a human stem cell-derived cardiomyocyte model. Journal of molecular and cellular cardiology 209, 143–160 (2025).

32. J. Zhao, L. Qi, S. Yuan, T. C. Irving, W. Ma, Differences in thick filament activation in fast rodent skeletal muscle and slow porcine cardiac muscle. The Journal of physiology 602, 2751–2762 (2024).

33. M. Reconditi, Recent Improvements in Small Angle X-Ray Diffraction for the Study of Muscle Physiology. Reports on progress in physics. Physical Society 69, 2709–2759 (2006).

34. M. Caremani et al., Dependence of myosin filament structure on intracellular calcium concentration in skeletal muscle. J Gen Physiol 155 (2023).

35. M. Reconditi et al., Thick Filament Length Changes in Muscle Have Both Elastic and Structural Components. Biophysical journal 116, 983–984 (2019).

36. M. Irving, Regulation of Contraction by the Thick Filaments in Skeletal Muscle. Biophysical journal 113, 2579–2594 (2017).

37. E. Brunello, L. Fusi, Regulating Striated Muscle Contraction: Through Thick and Thin. Annu Rev Physiol 10.1146/annurev-physiol-042222-022728 (2023).

38. D. Y. Barefield et al., Myosin-binding protein H-like regulates myosin-binding protein distribution and function in atrial cardiomyocytes. Proceedings of the National Academy of Sciences of the United States of America 120, e2314920120 (2023).

39. D. Dutta, V. Nguyen, K. S. Campbell, R. Padron, R. Craig, Cryo-EM structure of the human cardiac myosin filament. Nature 623, 853–862 (2023).

40. D. Tamborrini et al., Structure of the native myosin filament in the relaxed cardiac sarcomere. Nature 623, 863–871 (2023).

41. J. van der Velden et al., Isometric tension development and its calcium sensitivity in skinned myocyte-sized preparations from different regions of the human heart. Cardiovasc Res 42, 706–719 (1999).

42. T. Nakanishi et al., Effects of omecamtiv mecarbil on the contractile properties of skinned porcine left atrial and ventricular muscles. Front Physiol 13, 947206 (2022).

43. N. A. Narolska et al., Myosin heavy chain composition and the economy of contraction in healthy and diseased human myocardium. Journal of muscle research and cell motility 26, 39–48 (2005).

44. N. Piroddi et al., Tension generation and relaxation in single myofibrils from human atrial and ventricular myocardium. Pflugers Archiv : European journal of physiology 454, 63–73 (2007).

45. A. L. Hessel et al., The distinctive mechanical and structural signatures of residual force enhancement in myofibers. Proceedings of the National Academy of Sciences of the United States of America 121, e2413883121 (2024).

46. M. Reconditi et al., Myosin filament activation in the heart is tuned to the mechanical task. Proceedings of the National Academy of Sciences of the United States of America 114, 3240–3245 (2017).

47. E. Brunello et al., Myosin filament-based regulation of the dynamics of contraction in heart muscle. Proceedings of the National Academy of Sciences of the United States of America 117, 8177–8186 (2020).

48. P. M. Janssen, B. J. Biesiadecki, M. T. Ziolo, J. P. Davis, The Need for Speed: Mice, Men, and Myocardial Kinetic Reserve. Circulation research 119, 418–421 (2016).

49. J. H. Chung, B. J. Biesiadecki, M. T. Ziolo, J. P. Davis, P. M. Janssen, Myofilament Calcium Sensitivity: Role in Regulation of In vivo Cardiac Contraction and Relaxation. Front Physiol 7, 562 (2016).

