## Supplementary material for "Porcine Left Atrial and Ventricular Thick Filaments Exhibit Distinct Resting Structures and Calcium-dependent Responses": SI

**Short title: Atrial-Ventricular Differences in Thick Filament Structure and Calcium Responses**

Lin Qi^a,,1^, Maicon Landim-Vieira^a,1^, Hailey Flannagan^a^, Marcela Monroy^a^, Edward O. Olaniyan^a^, Meihua Guo^b^, Chengqian Gao^c^, Shengyao Yuan^a^, Henry Gong^a^, Suman Nag^d^, Thomas C. Irving^a,e,f^, ^f^, Weikang Ma^a,e,f,2^

a. Department of Biology, Illinois Institute of Technology, Chicago, IL 60616
b. Institute for Genome Engineered Animal Models of Human Diseases, National Center of Genetically Engineered Animal Models for International Research, Dalian Medical University, Dalian, Liaoning, 116044, China
c. College of Basic Medical Sciences, Dalian Medical University, Dalian, Liaoning, China
d. Department of Biochemistry, Bristol Myers Squibb (TM), Brisbane, CA 94005
e. Center for Synchrotron Radiation Research and Instrumentation, Illinois Institute of Technology, Chicago, IL 60616
f. Pritzker Institute of Biomedical Science and Engineering, Illinois Institute of Technology, Chicago, IL, USA

^1^L.Q. and M.L-V. contributed equally to this work.

**Results**

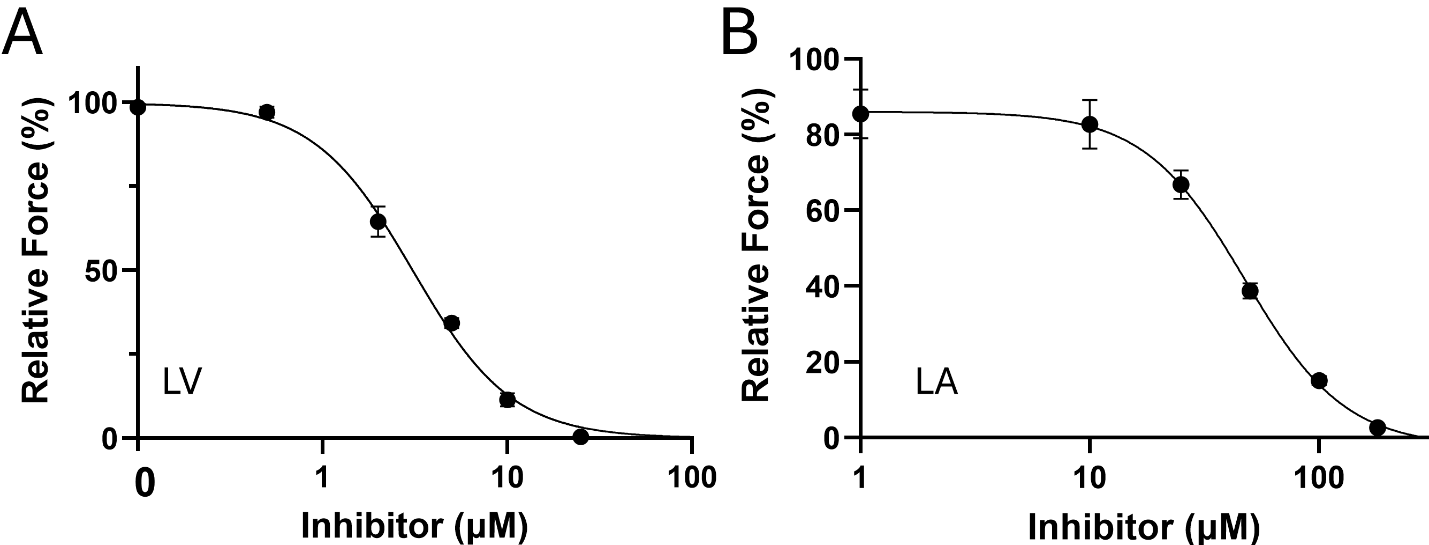

**Figure S1. Concentration-dependent relative maximum force**. The relative force from LV(**A**, replotted from (1)) and LA (**B**) in the increasing inhibitor (MYK-7600) concentrations.

The inhibitor suppressed force production in permeabilized porcine LA in a dose-dependent manner, with an IC50 of 47 μM at pCa 4.3 (Fig. S1B). This value is approximately one order of magnitude higher than the IC50 previously reported for LV at pCa 4.5 (3 μM) (1). At 180 μM inhibitor concentration, 2.6 ± 0.9% residual force remained. However, during the X-ray experiments, no detectable force development above baseline was observed in LA in the presence of the inhibitor at any pCa tested. This discrepancy may be explained by two factors. First, in the X-ray experiments, the LA tissues were incubated with 180 μM inhibitor for several hours before measurements, whereas the residual force observed in Fig. S1B was measured after only a 10 min incubation during the dose–response experiments. Second, the highest calcium concentration used in the dose–response experiments was pCa 4.3, corresponding to ~1.58-fold higher calcium concentration than the pCa 4.5 condition used in the X-ray experiments.

**TABLES**

**Table S1.**

|  |  | **LA** |  | **LV** |
| --- | --- | --- | --- | --- |
| **F_max (mN/mm_^2^_)_** |  | 61.04 ± 6.81** |  | 88.74 ± 5.10 |
| ***p*Ca_50_** |  | 5.89 ± 0.02 |  | 5.95 ± 0.04 |
| ***n*_Hill_** |  | 2.32 ± 0.20 |  | 2.39 ± 0.25 |
| ***k*_TR max_ (s^-1^)** |  | 13.56 ± 1.23*** |  | 7.48 ± 0.37 |
| **SS_max_ (MPa)** |  | 1.00 ± 0.22** |  | 2.06 ± 0.17 |
| **N** |  | 10 |  | 11 |

**Contractile parameters measured in porcine permeabilized left atria and ventricle.** F_max_, maximum steady-state isometric force; pCa_50_, Ca^2+^ concentration needed to reach 50% of the steady-state isometric maximum force; *n*_Hill_, cooperativity of thin filament activation; *k*_TR_ max, maximum rate of tension redevelopment; SS_max_, maximum steady-state sinusoidal stiffness; n, number of cardiac muscle preparations. Data are shown as mean ± SEM Statistical significance was assessed by unpaired Student's t-test. ** *p* < 0.01 and *** *p* < 0.001.

**Table S2.**

|  | SL (µm) | **3-state model fitted parameters** | | | | **3-state model predictions** | | |
| --- | --- | --- | --- | --- | --- | --- | --- | --- |
|  |  | Cross-bridge | | Regulatory unit | |  |  |  |
|  |  | *f* (s^-1^) | *g* (s^-1^) | *k*_ON_ (M^1^ s^1^) | *k*_OFF_ (s^1^) | pCa_50_ | max force (norm) | max *k*_TR_ (s^-1^) |
| **LA** | 2.3 | 6.16 | 7.81 | 1.84x10^8^ | 456.2 | 5.89 | 0.69 | 13.56 |
| **LV** | 2.3 | 4.87 | 2.77 | 1.84 x10^8^ | 581.7 | 5.95 | 1.00 | 7.48 |

**Optimized parameter predictions and estimates from the 3-state model for steady-state isometric force-*k*_TR_ data depicted in Fig. 1.** All best-fit parameter estimates for *f* (attachment rate), *g* (detachment rate), and *k*_OFF_ (calcium dissociation rate) were obtained in MatLab by using the simplex method for the steady-state isometric force-*k*_TR_ data shown in Fig. 1. The simplex method considers the values of myofilament calcium sensitivity (pCa_50_) from steady-state isometric force measurements as reported in Fig. 1; values for *k*_ON_ were derived from measurements reported by Pinto et al.(2). Maximum steady-state isometric force at LV was assumed to be 1.0. These parameter sets were used to fit and illustrate relations between *k*_TR_ and steady-state isometric force in Fig. 1G (dashed lines).

**References**

1.	W. Ma, S. Nag, H. Gong, L. Qi, T. Irving, Cardiac myosin filaments are directly regulated by calcium. *J Gen Physiol* **154** e202213213 (2022).

2.	J. R. Pinto *et al.*, Strong cross-bridges potentiate the Ca(2+) affinity changes produced by hypertrophic cardiomyopathy cardiac troponin C mutants in myofilaments: a fast kinetic approach. *The Journal of biological chemistry* **286**, 1005-1013 (2011).
